# The national alert-response strategy against cholera in Haiti: a four-year assessment of its implementation

**DOI:** 10.1101/259366

**Authors:** Stanislas Rebaudet, Gregory Bulit, Jean Gaudart, Edwige Michel, Pierre Gazin, Claudia Evers, Samuel Beaulieu, Aaron Aruna Abedi, Lindsay Osei, Robert Barrais, Katilla Pierre, Sandra Moore, Jacques Boncy, Paul Adrien, Edouard Beigbeder, Florence Duperval Guillaume, Renaud Piarroux

**Affiliations:** Assistance Publique – Hôpitaux de Marseille (AP-HM), Marseille, France; Hôpital Européen Marseille, Marseille, France; United Nations Children’s Fund, Haiti; Aix Marseille Univ, IRD, INSERM, SESSTIM, Marseille, France; Ministry of Public Health and Population, Directorate of Epidemiology Laboratory and Research, Haiti; Institut de Recherche pour le Développement (IRD), Marseille, France; Ministry of Health, Kinshasa, Democratic Republic of the Congo; Aix Marseille Univ, Marseille, France; Ministry of Public Health and Population, National Laboratory of Public Health, Haiti; Ministry of Public Health and Population, former Minister, Haiti; Sorbonne Université, INSERM, Institut Pierre-Louis d’Epidémiologie et de Santé Publique, AP-HP, Hôpital Pitié-Salpêtrière, F-75013, Paris, France

## Abstract

**Background:** A massive cholera epidemic struck Haiti on October 2010. As part of the national cholera elimination plan, the Haitian government, UNICEF and other international partners launched a nationwide alert-response strategy from July 2013. This strategy established a coordinated methodology to rapidly target cholera-affected communities with WaSH (water sanitation and hygiene) response interventions conducted by field mobile teams. An innovative *red-orange-green* alert system was established, based on routine surveillance data, to weekly monitor the epidemic.

**Methodology/Principal findings:** We used cholera consolidated surveillance databases, alert records and details of 31,306 response interventions notified by WaSH mobile teams to describe and assess the implementation of this approach between July 2013 and June 2017. Response to *red* and *orange* alerts was heterogeneous across the country, but significantly improved throughout the study period so that 75% of *red* and *orange* alerts were responded within the same epidemiological week during the 1^st^ semester of 2017. Numbers of persons educated about cholera, houses decontaminated by chlorine spraying, households which received water chlorination tablets and water sources that were chlorinated during the same week as cholera alerts significantly increased. Alerts appeared to be an interesting and simple indicator to monitor the dynamic of the epidemic and assess the implementation of response activities.

**Conclusions/Significance:** The implementation of a nationwide alert-response strategy against cholera in Haiti was feasible albeit with certain obstacles. Its cost was less than USD 8 million per year. Continuing this strategy seems essential to eventually defeat cholera in Haiti while ambitious long-term water and sanitation projects are conducted in vulnerable areas. It constitutes a core element of the current national plan for cholera elimination of the Haitian Government.

## Introduction

On October 2010, cholera was inadvertently introduced in Haiti [1], the poorest country of the Western Hemisphere [2]. It then experienced the largest national-level epidemic of the past few decades with 816,066 suspected cases and 9,748 suspected deaths recorded on December 30, 2017 by the Haitian Ministry of Public Health and Population (MSPP, acronyms summarized in Supplementary Table S1) [3]. Following international recommendations [4], a cholera surveillance and early warning system was established while Haitian authorities and services, together with numerous international and nongovernmental organizations (NGOs), struggled to mitigate the death toll and case incidence by supporting cholera treatment institutions and providing efforts to provide safe water, and to improve sanitation and hygiene practices (WaSH) in affected communities [5–8]. Cholera incidence gradually receded in 2011–2012, with alternating troughs and peaks influenced by seasonal rainfall [9]. Although Haiti then remained the most cholera-affected country worldwide [10], emergency funds for cholera dried up and most organizations eventually interrupted or drastically reduced their activities in 2012 [7,11]. The few remaining programs often lacked the precise epidemiological data necessary to target their efforts to the appropriate areas [11]. On February 2013, the Haitian government, Pan American Health Organization (PAHO), United Nations Children’s Fund (UNICEF) and the Centers for Disease Control and Prevention (CDC) launched an ambitious National Plan for the Elimination of Cholera in Haiti 2013–2022 [12]. Over USD 1.5 billion of the total USD 2.2 billion was designated to invest in Haitian water and sanitation infrastructures, while only 68% of households drank from improved water sources, 26% had access to improved sanitation facilities and 34% had water and soap available for hand washing [13]. Meanwhile, two pilot oral cholera vaccination campaigns vaccinated approximately 100,000 people in both rural and urban settings [14,15], and two additional campaigns were planned in 2013 [16].

The elimination plan also intended to improve surveillance activities and ensure adequate response to detected outbreaks [12]. In order to interrupt cholera local outbreaks at an early stage, UNICEF thus backed the MSPP and the Haitian National Directorate for Water and Sanitation (DINEPA) to launch a cholera alert-response strategy in July 2013. This program aimed to rapidly detect cholera local outbreaks and send response teams to affected communities in order to identify additional cases, to educate on risk factors, prevention and management methods, to distribute soap and water chlorination products, and to protect local water sources by establishing water chlorination points. At central level, the epidemic was monitored using a simple and original cholera alert system that classified each commune in *red, orange* or *green* on a weekly basis, according to standardized criteria.

Based on identified risk factors, and on the growing evidence of WaSH efficiency against cholera [17–20] or against diarrhea in developing countries or humanitarian crises [21–24], promoting hygiene and improving access to safe drinking water have for long been recommended to control cholera transmission [25,26,4,27]. Besides, case-area targeted interventions have been frequently conducted against cholera epidemics, and are supported by the observed increase of cholera risk for neighbors living within a few dozen meters of cases during days following their presentation [28,29]. However, feedback from the field has been scarce and scattered, reported activities have usually been implemented at a local level, during short time periods, and described with few details [30–32,26,33–38,7,8,39–43].

To our knowledge, we thus propose in the present study the first thorough and quantified description of a multiyear nationwide coordinated strategy aiming to mitigate cholera transmission at the community level through rapid and targeted WaSH response interventions. Implementation of the National alert-response strategy against cholera in Haiti was assessed between July 2013 (27th epidemiological week, 2013w27) and June 2017 (2017w26). Evaluation of the efficiency and impact of this strategy is out of this paper’s scope.

## Methods

### Study site

The Republic of Haiti occupies the mountainous western third of the island of Hispaniola, in the Greater Antilles. The country spans over 27,000 km^2^, and the estimated population in 2015 was approximately 10.9 million, 52% of whom lived in urban areas [44]. Haiti is administratively composed of 10 departments and 140 communes, which range in surface area from 59 to 645 km^2^ and host populations between less than 6,000 and over 987,000 inhabitants [44]. Economic collapse, demographic explosion, political instability and natural disasters have contributed to generate deep structural vulnerabilities in WaSH, health care, education, agriculture, environment, trade, transportation and governance sectors that may favor cholera transmission [45].

### The cholera surveillance system in Haiti

Since October 2010, cholera treatment institutions (between 136 and 262 cholera treatment centers [CTCs], units [CTUs], or acute diarrhea treatment centers [CTDAs] during the study period) routinely record and notify cholera-associated morbidity and mortality data. According to the WHO standard definition [4], a probable cholera case is defined as a patient aged 5 years or older who develops acute watery diarrhea, with or without vomiting. In Haiti, suspected cholera cases aged <5 years old with similar symptoms are also separately recorded and included in the global cholera toll. Daily seen and hospitalized suspected cases as well as daily institutional and community suspected deaths are anonymously transmitted to one of the ten department health directorates using formatted SMSs (Short Message Service) or phone calls (Supplementary Figure S1). Department directorates compile and validate this data using a Microsoft Excel spreadsheet, which is sent to the national Directorate for epidemiology laboratory and research (DELR), normally on a daily basis, but with frequent delays. The DELR validates the departmental data and compiles them into a national Microsoft Excel spreadsheet at a daily and communal scale, before performing analyses. Recorded communes refer to the treatment place of cholera patients, not to their residence. Community cases are not recorded.

Since 2010, routine bacteriological confirmation of suspected cholera cases is performed at the National Laboratory of Public Health (LNSP) in Port-au-Prince Metropolitan Area, using standard sampling, culture and phenotyping methods [46]. Since July 2017, a private lab run by the NGO Zanmi Lasante and located in Saint-Marc, Artibonite, notifies stool culture results to LNSP. By June 2017, a total of 13,625 stool specimens, mostly sampled using rectal swabs with Carry-Blair transport medium, had been included in the national microbiological surveillance system of cholera. Sampling is not homogeneous across the country and about 1/3 of specimens actually have originated form a sentinel surveillance network of four hospitals in three departments [47]. The remainder has been sampled in other CTCs/CTUs/CTDAs across the country by the staff of department health directorates as well as medical international or non-governmental organizations. All partners have been committed to transport specimens back to the LNSP in Port-au-Prince.

Between 2013 and 2015, the MSPP experimented with the nationwide use of rapid diagnostic tests (Crystal VC^TM^ RDT, Span Diagnostics Ltd., India) in cholera treatment institutions. Training proved challenging, and RDT results were barely taken into account for routine surveillance, alert identification and response interventions as initially planned.

### The nationwide response strategy to cholera suspect cases

The National alert-response strategy against cholera in Haiti was launched on July 2013. It planned to improve: (a) the coordination of activities implemented by national, international and nongovernmental partners involved in cholera control; (b) the epidemiologic surveillance of cholera in every commune, and the monitoring of outbreaks via an alert detection system at the central level; (c) the rapidity, exhaustiveness, targeting and relevance of field responses to cholera alerts; and, (d) cholera prevention in the most vulnerable areas using mass education sessions and rehabilitation or installation of water adduction infrastructures. For this, UNICEF contracted at least one WaSH international or nongovernmental organization for each of the 10 departments (Supplementary Table S1), which hired rapid response mobile teams composed of local Haitian staff. These WaSH teams were requested to address every cholera suspected case or death by an intervention in the neighboring community within 48 hours. In case of several concomitant outbreaks, they were asked to target most affected areas in priority. The core methodology of their response interventions, established with the Ministry of Health and its partners [48], involved: (i) verification of surveillance data in register books of treatment institutions and the identification of affected localities and neighborhoods; (ii) field investigations in affected communities to estimate the extent of cholera transmission, understand triggering/aggravating factors, and to identify contacts and suspected cases with the help of community leaders and community officers; (iii) visits to affected families and their neighbors (minimum 5 households depending on the local geography), who were proposed house decontamination by chlorine spraying, although efficacy and impact of this decontamination method has never been established [27] and is likely limited for a few hours; (iv) on-site organization of education sessions about cholera transmission modes, prevention and initial care methods; (iv) distribution of 1 cholera kit per household (composed of 5 soaps, 5 sachets of oral rehydration salts (ORS), and about 115 chlorine tablets – 80 Aquatabs^TM^ 33mg in urban or 150 in rural areas); (v) set up of manual bucket chlorination at drinking water sources during 1 or more weeks when possible, by hiring and instructing local volunteers; (vi) repair and extra-chlorination of water adduction systems when necessary and possible. Activities implemented during response interventions to cholera cases were prospectively transmitted by WaSH mobile teams to UNICEF, using standardized online Google spreadsheets. A few other WaSH organizations implementing field response to cholera cases funded by other agencies decided to join the strategy and also reported their activities to UNICEF (Supplementary Table S1).

To bolster institutional response capacities, UNICEF and the World Bank also provided additional material, funds and human resources to DINEPA and to MSPP which created its own departmental response teams (EMIRAs, *Rapid intervention mobile teams*) on March 2014, with WaSH and medical personnel (Supplementary Table S1). Besides activities listed above, their terms of reference included: (vii) primary care of cholera cases found in the community; (viii) chemoprophylaxis of contacts living in the same house as cholera cases with one dose of doxycycline 300 mg for non-pregnant adults only [4,49,50]; (ix) nursing support to cholera treatment institutions when necessary. ECHO (European Commission Humanitarian Office) and PAHO (Pan American Health Organization) also contracted medical NGOs (Supplementary Table S1), with similar terms of reference as EMIRAs’. Reporting of EMIRAs’ and medical mobile teams’ activities could not be systematized as for WaSH teams contracted by UNICEF.

The medical and WaSH governmental and nongovernmental actors of each department were requested to organize at least monthly coordination meetings and to share cholera epidemiological data and rumors on a daily basis (Supplementary Figure S1). Field interventions integrating WaSH/medical and governmental/nongovernmental staff were strongly encouraged. Mobile teams were asked to repeat response interventions in the community until every suspected cholera case had been addressed. From 2017, they were also requested to perform post distribution monitoring two weeks later. An illustration of these response interventions can be seen on this short online video (https://www.youtube.com/watch?v=KOYRX4Fmabo, accessed 26 January 2018).

### The cholera alert system

To better monitor the epidemic and check that field actors targeted probable cholera transmission foci, epidemiologists and WaSH specialists from the DELR, Assistance Publique - Hôpitaux de Marseille (AP-HM) and UNICEF established a simple alert system based on cholera surveillance data at the communal scale in July 2013. Three stratified levels of alert – *red, orange* and *green* alert – were defined based on criteria including the number of cholera suspected cases and associated deaths (≥ 5 years of age only) during the past seven days (Table 1). This approach was consensually and officially validated on August 2013 during a national Cholera Alert and Response workshop with Haitian authorities and international partners. Filling routinely compiled surveillance data in a Microsoft Excel programmed spreadsheet, DELR staff prospectively computed weekly alerts. But for organizational reasons, they actually did not include all planned criteria (Table 1), such as precise location of cholera cases at a sub-communal scale (criteria #3), RDT results (#4), stool cholera cultures (#5), rumors of cholera outbreak (#9). DELR then mapped identified alerts using Philcarto software [51], and summarized alert information in cholera bulletins, which were diffused to partners via epidemiological situation rooms, emails or through the MSPP website, usually on a weekly basis. An example of cholera weekly bulletin can be seen online (https://mspp.gouv.ht/site/downloads/Profil%20statistique%20Cholera%2044eme%20SE%202016%20.pdf, accessed 26 January 2018).

**Table 1.** – Criteria for cholera alerts at the communal level according to the DELR*

|  |  |  |
| --- | --- | --- |
| **Red alert** |  |  |
| #1<sup>SE</sup> | | $\geq 1$ cholera-associated hospital or community death of individual ≥ five years of age during the past seven days of record |
| #2<sup>SE</sup> | and/or | $\geq 10$ suspected cholera cases aged ≥ five years during the past seven days of record |
| #3 | and/or | $\geq 5$ suspected cholera cases aged ≥ five years originating from a similar locality during the past seven days of record |
| #4 | and/or | $\geq 50\%$ of positive rapid diagnostic tests (RDT) |
| #5<sup>E</sup> | and/or | $\geq 1$ stool culture positive for *Vibrio cholerae* O1 at the LNSP* |
| **Orange alert** |  |  |
| #6<sup>SE</sup> |  | No red alert criteria |
| #7<sup>SE</sup> | and | Twofold or more increase in suspected cases aged ≥ five years during the past seven days compared with the previous seven-day period |
| #8<sup>SE</sup> | and/or | Red alert during the previous week |
| #9 | and/or | Outbreak rumor<sup>S</sup> in the commune |
| **Green alert** |  |  |
| #10<sup>SE</sup> |  | No red or orange alert criteria for at least two weeks |

\* DELR, Haiti Directorate for Epidemiology Laboratory and Research; LNSP, National Laboratory of Public Health
<sup>S</sup> Criteria that were prospectively used by the DELR on a weekly basis (#1 #2 #6 #7 #8 #10) using often incomplete and delayed cholera surveillance reports; the prospective collection and analysis of data actually appeared to be difficult for the other criteria
<sup>E</sup> Criteria that were retrospectively used in the study to identify retrospective alerts (#1 #2 #5 #6 #7 #8 #10), using consolidated databases.
<sup>S</sup> Rumors define any information about cholera outbreaks that is not produced by the surveillance system of suspected cases and deaths and stool cultures

### Data collection

We used anonymous information of daily reported cholera suspected cases, deaths and stool culture results, with the authorization of, and in collaboration with, the Haitian Ministry of Health authorities. The dates of alert identification by the DELR and the dates of diffusion to partners via alert bulletin email were prospectively collected. UNICEF provided the list of its expenditure for mobile teams (through international or non-governmental organizations, and MSPP with EMIRAs) and for other cholera related activities such as cholera surveillance, coordination of partners and other WaSH prevention activities. UNICEF also provided a list of response items (chlorine, soaps, buckets, oral rehydration salts…) delivered to its partners for response and prevention activities, as well as their cost. Records and details of field activities notified by WaSH mobile teams included the date, location (*i.e.* commune, communal section, locality) and nature (*i.e.* investigation, education sessions, distribution of soap and chlorine, implementation of water-chlorination points, household decontamination…). For each response, field organizations mentioned the main institution that notified related cases, so that it was possible to link field activities with institutional epidemiological information. Response data were used with the consent of health authorities, implementing organizations and funders. Country and commune population estimates were provided by the Haitian Institute of Statistics and Informatics (http://www.ihsi.ht/pdf/projection/Estimat_PopTotal_18ans_Menag2015.pdf accessed 26 January 2018). Satellite estimates of daily-accumulated rainfall (area-averaged TRMM_3B42_daily v7) were extracted from NOAA websites on the entire surface of Haiti, and on the centroid of each of the 140 communes (http://giovanni.gsfc.nasa.gov/giovanni/, accessed 26 January 2018). Corresponding databases are listed in Supplementary Table S2.

### Retrospective identification of cholera alerts

Because the cholera alert system was launched to prospectively monitor the epidemic, we used cholera alerts as an outbreak proxy to retrospectively evaluate the implementation of the alert-response strategy. Hence, we could detect delays in notification and analysis of cholera routine surveillance data at DELR, as well as the capacity of response teams to directly get epidemiological information from the community, treatment institutions and departmental directorates of health. We computed retrospective cholera alerts classified as *red, orange* or *green* at commune-level and per epidemiological week, between July 2013 (27th epidemiological week, 2013w27) and June 2017 (2017w26), using consolidated databases of institutional cholera suspected cases, suspected deaths and of stool culture results, as well as alert criteria including results of stool culture for cholera (criteria #5) (Table 1). Retrospective alerts were then used as a proxy to: (i) describe the epidemic; (ii) assess the surveillance system by comparing alerts that DELR prospectively identified and communicated at the central level with alerts that *should* have been prospectively identified and communicated; and, (iii) assess the field response implemented by mobile teams to outbreaks that *should* have been responded.

Weekly evolution of alerts was plotted at national levels. The relative proportion *of red, orange* and *green* cumulated alerts was mapped at the communal level. Main alert characteristics (commune population, accumulated rainfall in the commune, number of suspected cases and deaths, number of stool samples received at the LNSP lab for cholera confirmation and culture positivity ratio) were summarized for each alert level and plotted for each alert level, each of the 8 semesters of the 4-year study period, each of the 10 departments.

### Evaluation of the implementation of the alert-response strategy

To evaluate the implementation of the alert-response strategy, we linked retrospective alerts with alerts actually identified and communicated by DELR and with response intervention conducted in the same commune. We then assessed the proportions of *red and orange* retrospective alerts, which were (1) identified by the DELR, (2) communicated by email or on the MSPP website, and, (3) responded by WaSH mobile teams during the same epidemiological week and during the same or following week, considering that outbreaks may start on weekends. We also counted the number of WaSH response interventions conducted for each alert. Each indicator was plotted by alert level, semester and department. The differences between alert levels (*red* versus *orange*), the evolution along the semesters of the study (from 1 to 8) and the heterogeneity between the 10 departments were estimated via multivariate analysis using generalized linear mixed models [52]. This approach allowed us to take into account two levels of spatial heterogeneity, by modeling departments and communes as nested random effect variables. Using a binomial distribution for alert identification, communication and response rates, models estimated odds ratios for alert levels and semesters, and *P*-values. Using a negative binomial distribution for numbers of response interventions, of educated persons, of decontaminated houses, of households which received chlorine tablets and of chlorinated water sources, models estimated relative risks and *P*-values.

To assess the capacity of WaSH mobile teams to independently identify and respond to cholera cases, we looked at the alert level communicated by DELR for communes where interventions were conducted and plotted these counts by semester and department. For each intervention, we also counted and plotted the number of cases notified in the commune during the same week. The numbers of people targeted by education sessions, of houses decontaminated by chlorine spraying, of households who received chlorine tablets, and of water sources that were chlorinated was computed for each response intervention, and plotted by semester and department.

### Software and packages

Data management was performed using Microsoft^®^ Excel for Mac v15.32. The map was drawn using QGIS v2.18 (http://www.qgis.org, accessed 26 January 2018). Graph design and statistical analyses were performed using RStudio version 1.0.136 for Mac (http://www.rstudio.com/, accessed 26 January 2018) with R version 3.4.2 for Mac (http://www.r-project.org/, accessed 26 January 2018) and the ggplot2 [53] (https://cran.r-project.org/web/packages/ggplot2/index.html, accessed 26 January 2018), and lme4 [54] (https://cran.r-project.org/web/packages/lme4/index.html, accessed 26 January 2018) packages.

## Results

### Evolution of the epidemic and timeframe of the response strategy

Between the launch of the nationwide alert-response strategy in July 2013 (2013w27) and the end of this 209-week study period in June 2017 (2017w26), 149,690 suspected cases and 1,498 deaths were notified across Haiti (Figure 1 Panel A). A total of 8,094 stool samples were cultured and 52% of them were positive for *V. cholerae* O1 (Figure 1 Panel A).

**Figure 1.**
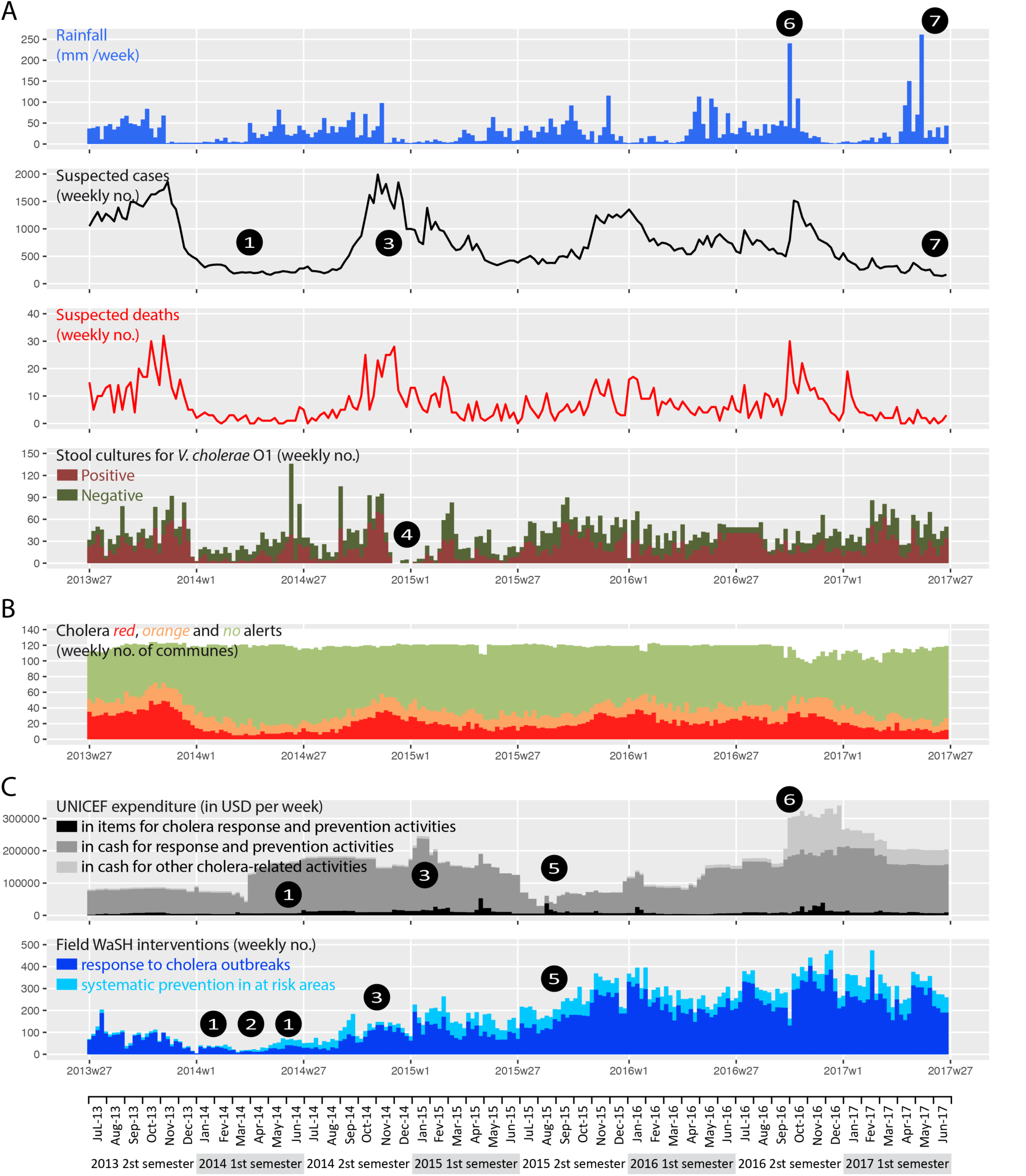
– Weekly evolution of: accumulated rainfall and cholera epidemic indicators (Panel A); cholera retrospective alerts (Panel B); and implementation of the response strategy by UNICEF (Panel C), from mid-2013 (2013w27) to mid-2017 (2017w26) Area-averaged cumulated rainfall was obtained from NOAA. Suspected cholera cases, cholera-associated deaths, positive and negative stool cultures tested for *V. cholerae O1* were provided by routine surveillance databases of the MSPP (Ministry of Public Health and Population). Details on expenditure and field WaSH (Water Sanitation and Hygiene) interventions were provided by UNICEF. *Response* refers to activities (i) to (vi) described in the Methods section, implemented in response to cholera cases. *Prevention* refers to activities (i) to (vi) implemented systematically in at risk areas. *Other cholera-related activities* refers to cholera surveillance, coordination of partners, as well as other WaSH prevention activities *(i.e.* in response to hurricane Matthew **❻** from October 2016). **❶**, prolonged low incidence period in 2014, despite regular precipitation from March and few WaSH response interventions regardless of the increase of funds provided by UNICEF to partner organizations from April; the temporary drop in March was consecutive to the Chikungunya epidemics and a halt of contracts with several partner organizations; **❷**, launch of EMIRAs by MSPP from March 2014. **❸**, cholera outbreaks in the Port-au-Prince Metropolitan Area from September 2014 involving vandalism of main water pipes by gangs; escalation of interventions, and peak of funding in January 2015 in response to the crisis. **❹**, no stool cultures could be performed between December 2014 and January 2015 because of a shortage of laboratory reagents at the National Public Health Lab (LNSP). **❺**, intensification of the response strategy despite a contraction of funding. **❼**, cholera incidence decrease in spite of important rainfall.

During the same period, UNICEF expended USD 25.4 million through international or nongovernmental organizations for WaSH mobile teams and with MSPP for EMIRAs, USD 2.0 million in response items (chlorine, soaps, buckets, oral rehydration salts…), and an additional USD 3.7 million for other cholera related activities such as cholera surveillance, coordination of partners and other WaSH prevention activities (Figure 1 Panel C). UNICEF delivered 3.3 million soaps, 140 million AQUATABS^TM^ 33mg tablets and 3.6 million oral rehydration salts (ORS) sachets to UNICEF partner organizations (data not shown). A total of 31,306 interventions in response to cholera cases and 9,540 systematic prevention interventions were notified by WaSH partners of UNICEF (Fig 1 Panel C). Their mobile teams provided education sessions to 2.9 million people, decontaminated 179,830 houses, distributed chlorine tablets to 757,693 households as well as soaps to 593,494 households, and supplied chlorination at 2,282 water sources or networks. Unfortunately, field interventions of EMIRAs and mobile teams of medical international or non-governmental organizations could not be exhaustively counted and described, although most WaSH interventions integrated medical and governmental staff. Notably, information concerning the use of doxycycline chemoprophylaxis was not available for analysis.

The strategy started during the 2013 rainy season, and the case load slowly rose to 1,614 weekly suspected cases in November 2013 (Figure 1). Cholera incidence then collapsed, concomitantly with the dry season and around 50 weekly response interventions. An unprecedented 38-week period with under 500 notified cases weekly (including 23 weeks under 250 cases) expended until late September 2014, despite regular precipitation from March and few notified field interventions regardless of the increase of available funds. An abrupt increase in cholera incidence to 1,990 at 2014w45 was then observed from September, which mainly affected the Port-au-Prince Metropolitan Area and notably involved vandalism of main water pipes by gangs (results from unpublished field investigations). This crisis in the capital persisted throughout the 2014–2015 dry season, and stimulated a marked intensification of field interventions by WaSH partners. A second step in intensification of the strategy occurred during 2015 2^nd^ semester despite a notable but temporary reduction of available funds. Years 2015 and 2016 exhibited a sustained cholera case load oscillating between 500 and 1,500 per week with seasonal fluctuations, notably following hurricane Matthew that stroke South and Grand’Anse departments in October 2016. Additional funds were obtained so that a high volume of field interventions could be maintained until the end of the study period in June 2017. Concomitantly, cholera incidence exhibited a continuous decrease and remained below 500 weekly cases in 2017, although exceptional precipitations were recorded in April and May (Figure 1).

### Time-space distribution and description of cholera retrospective alerts

Using completed cholera databases of case, death and stool culture, and, the DELR criteria (Table 1), we retrospectively computed a total of 4,378, 3,475 and 16,710 *red, orange* and *green* alerts, respectively, across all 140 communes and during the 209 weeks between mid-2013 (2013w27) and mid-2017 (2017w26). A median weekly number of 20 communes were in *red alert* (Table 2). But as expected, alerts exhibited a temporal evolution consistent with the dynamic of the epidemic (Figure 1 Panel B), with a weekly minimum of 5 *red* communes observed in March-April 2014, and a maximum of 49 in October-November 2013. The distribution of cholera alerts also showed a marked spatial heterogeneity, as *red* and *orange* alerts mainly clustered in the departments of Ouest (DSO, especially in Port-au-Prince Metropolitan Area), Centre (DSC) and Artibonite (DSA) (Figure 2).

**Table 2.** – Characteristics of retrospectively identified *red, orange* and *green* alerts

|  | All alert levels | *Red* | *Orange* | *Green* |
| --- | --- | --- | --- | --- |
| No. of commune-weeks | 24563 | 4378 | 3475 | 16710 |
| (% of total) |  | (18%) | (14%) | (68%) |
| Median weekly no. of communes in alert |  | 20 | 16 | 80 |
| [min-max] |  | [5-49] | [6-28] | [45-107] |
| Median population of the commune, x1000 inhabitants | 40.5 | 119 | 48.3 | 34.1 |
| [min-max] | [5.7-972] | [7.6-972] | [7.6-972] | [5.7-972] |
| Median weekly accumulated rainfall in the commune, mm | 9 | 15 | 9 | 8 |
| [min-max] | [0-396] | [0-385] | [0-396] | [0-396] |
| Median no. of suspected cases per alert | 0 | 19 | 3 | 0 |
| [min-max] | [0-698] | [0-698] | [0-21] | [0-24] |
| Median no. of suspected deaths per alert | 0 | 0 | 0 | 0 |
| [min-max] | [0-13] | [0-13] | [0-2] | [0-2] |
| No. of stool samples received at the LNSP for *V. cholerae* culture | 8379 | 7595 | 325 | 459 |
| Culture positivity ratio, % | 53% | 58% | 0% | 0% |

**Figure 2.**
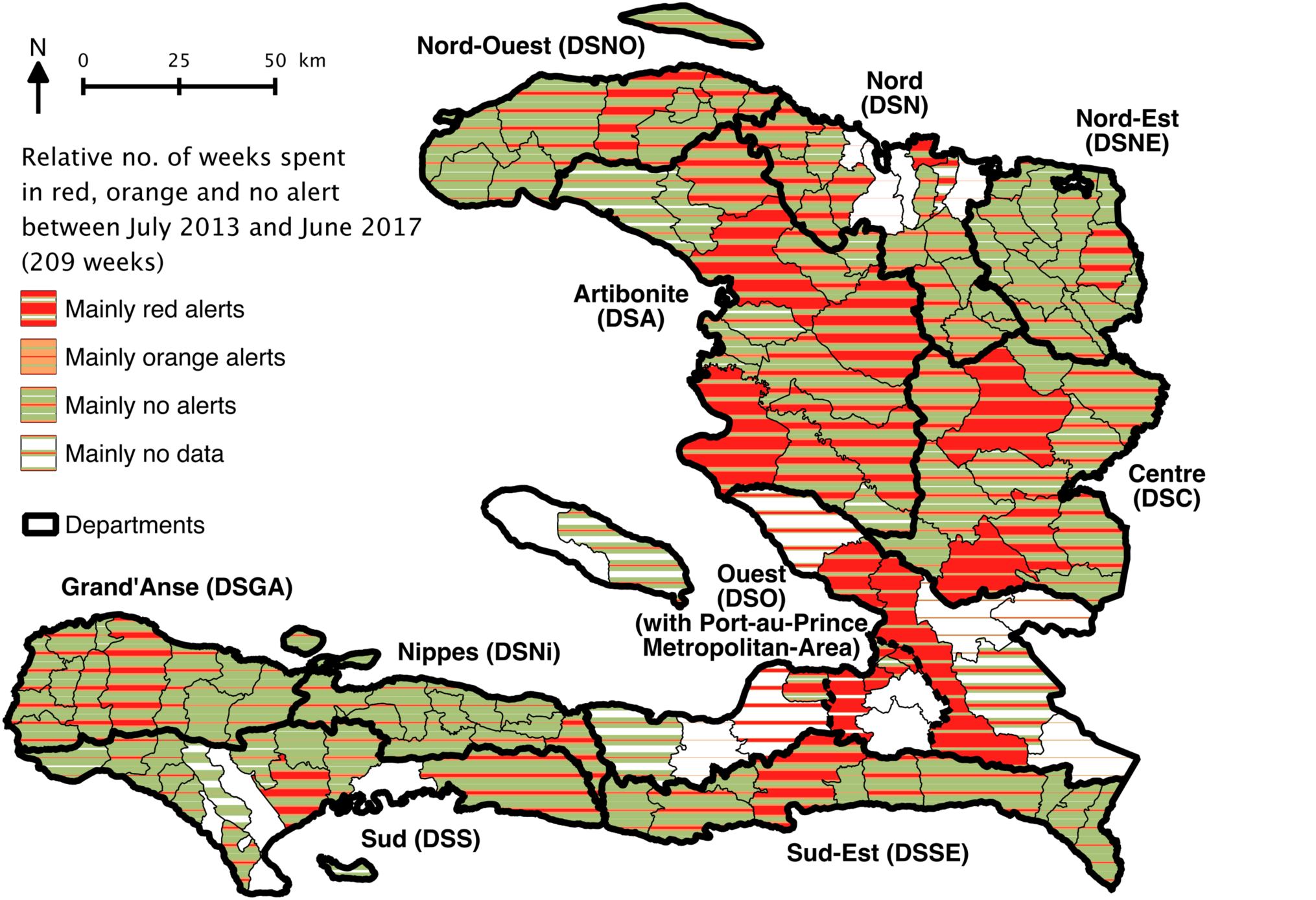
– Communal distribution of retrospectively identified alerts from July 2013 (2013w27) to June 2017 (2017w26). The respective number of weeks that each commune spent in red, orange or green alert, is indicated via a stripe pattern gradient.

Rainfall accumulated during *Red* alerts appeared more important than during *orange* alerts or in the absence of alert (*green*) (Table 2, Supplementary Figure S2). *Red* alerts were generally located in more populated communes as compared with *orange* or *green* alerts, but they affected from very small rural to highly populated urban communes (Table 2, Supplementary Figure S2). Additionally, *red* alerts corresponded to a variety of epidemic situations (Table 2): some of which were associated with less than 10 weekly cases and no reported deaths, but a single positive *V. cholerae* O1 culture that suggested low-grade confirmed cholera transmission, while other such alerts corresponded to more than 100 suspected cases with several associated deaths, or numerous stool samples all positive for *V. cholerae* O1 (Table 2, Supplementary Figure S2).

### Identification and communication of alerts by DELR

From July 2013 to June 2017, DELR staff performed cholera alert analyses for 192 of the 209 studied epidemiological weeks (92%). To identify cholera alerts, DELR actually never used patient addresses, rapid diagnostic test results, stool culture results or outbreak rumors, as initially proposed (Table 1), because prospective collection and analysis of these data appeared to be difficult. Via weekly epidemiological situation rooms, emails or through the MSPP website, DELR communicated alert analyses covering 181 of the 209 studied epidemiological weeks (88%). But alert analyses were not systematically complete.

Using retrospective alerts computed on the 2013w27 – 2017w26 period as the standard, DELR correctly identified 48% of *red* and *orange* alerts during the same week, 60% during the same or the following week (Table 3). DELR communicated 35% of *red* and *orange* alerts during the same week and 57% during the same or the following week (Table 3). Using generalized linear mixed models taking into account the time evolution (semesters along the study) and the heterogeneity between departments, alert identification and communication appeared significantly better for *red* than for *orange* alerts, whether during the same week (respective odds ratios (ORs) [95%-CI], 6.05 [5.39–6.80] and 8.61 [7.50–9.91]) or during the same or following week (Figure 3 Panel A&B, Table 3). Along the course of the study period, identification of *red* and *orange* alerts significantly improved during the same week but not during the same or following week (Figure 3 Panel A, Table 3). Conversely, communication during the same week remained stable, but improved during the same or following week (Figure 3 Panel B, Table 3). In particular, alert identification and communication dramatically dropped during the first semester of 2017 (Figure 3 Panel A&B). Identification and communication exhibited a significant heterogeneity between the 10 departments (*P*-values < 0.0001) (Figure 3 Panel A&B, Table 3).

**Table 3.** – Identification, communication of cholera alerts by DELR, and response to cholera alerts by WaSH mobile teams during the same week and during the same or following week, from July 2013 to June 2017: difference between alert levels, evolution along the eight semesters of the study period, and heterogeneity between departments.

| *Red or orange alerts*<br>(n = 7,853): | no<br>(%) | Difference<br>between alert levels<br>(*Red* vs *Orange* alerts) |  | Evolution<br>throughout the study<br>(per semester since July<br>2013) |  | Difference between<br>departments<br>(vs DSO*) |
| --- | --- | --- | --- | --- | --- | --- |
|  |  | Odds ratio<br>[95%-CI] | P-value | Odds ratio<br>[95%-CI] | P-value | P-value |
| Identification by DELR* | 3,800 | 6.05 | < 0.0001 | 1.14 | < 0.0001 | < 0.0001 |
| during the same week | (48%) | [5.39-6.80] |  | [1.12-1.17] |  |  |
| Identification by DELR* | 4,710 | 6.23 | < 0.0001 | 1.02 | 0.17 | < 0.0001 |
| during the same or following week | (60%) | [5.54-7.02] |  | [0.99-1.04] |  |  |
| Communication by DELR* | 2,725 | 8.61 | < 0.0001 | 1.01 | 0.40 | < 0.0001 |
| during the same week | (35%) | [7.50-9.91] |  | [0.99-1.04] |  |  |
| Communication by DELR* | 4,509 | 5.86 | < 0.0001 | 1.11 | < 0.0001 | < 0.0001 |
| during the same or following week | (57%) | [5.22-6.58] |  | [1.08-1.13] |  |  |
| Response by WaSH mobile teams | 3,822 | 2.52 | < 0.0001 | 1.62 | < 0.0001 | < 0.0001 |
| during the same week | (49%) | [2.23-2.86] |  | [1.57-1.66] |  |  |
| Response by WaSH mobile teams | 4,604 | 2.52 | < 0.0001 | 1.68 | < 0.0001 | < 0.0001 |
| during the same or following week | (59%) | [2.22-2.87] |  | [1.63-1.73] |  |  |

\* DELR, Directorate of Epidemiology Laboratory and Research of the Ministry of Health (MSPP); DSO, West department

**Figure 3.**
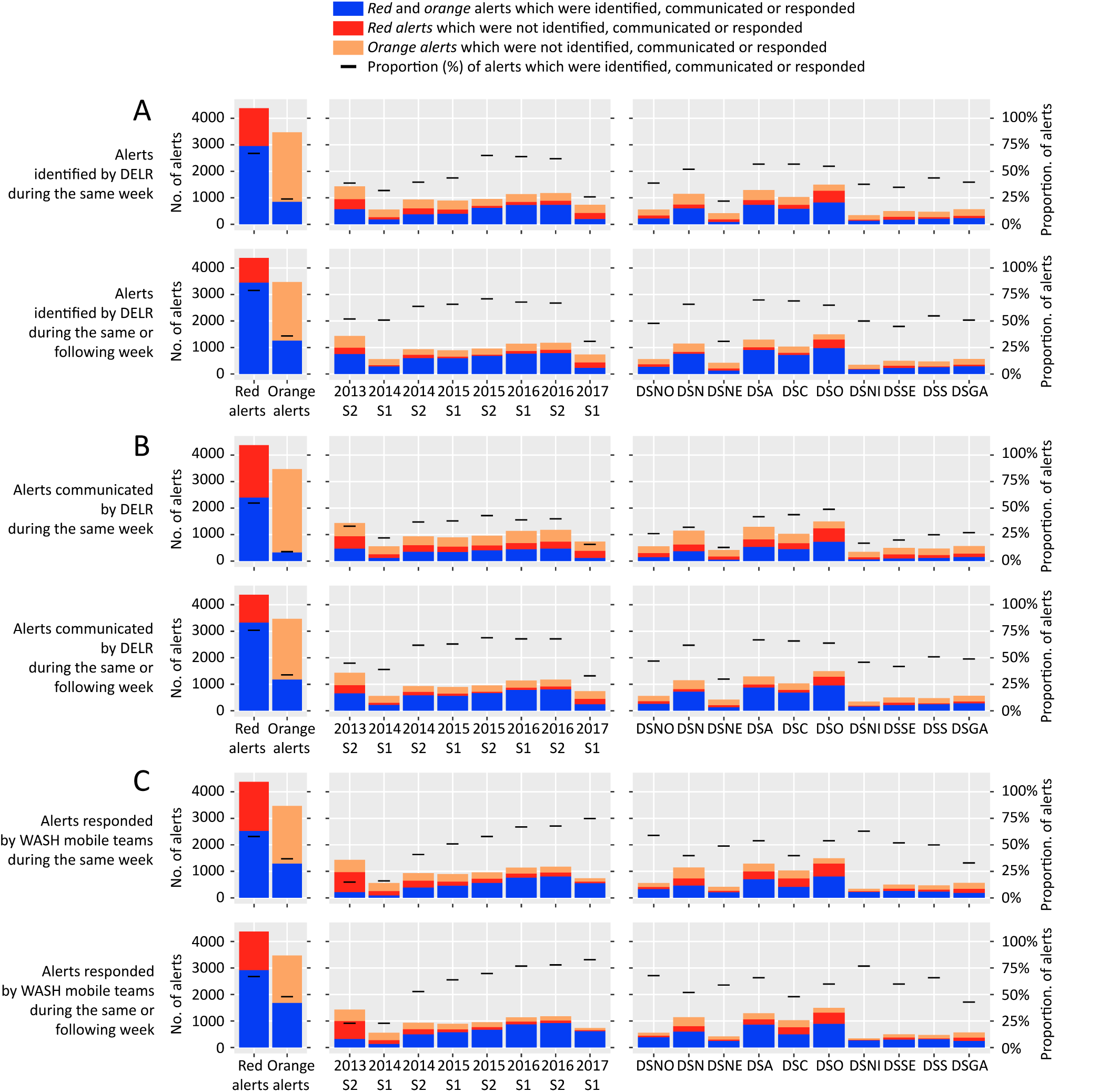
– Identification of cholera alerts by DELR (Panel A), communication of cholera alerts by DELR (Panel B) and response to cholera alerts by WaSH mobile teams (Panel C), during the same week and during the same or following week, from July 2013 to June 2017: difference between alert levels (*red* vs *orange*), evolution along the eight semesters of the study period, and difference between departments. Stacked bar plots show the following: in blue, the number of *red* or *orange* alerts which were identified, communicated or responded during the same week, and during the same or following week; in red and in orange, the number of *red* or *orange* alerts which were *not* identified, communicated or responded during the same week, and during the same or following week. Black dashes show the proportion of *red* or *orange* alerts which were identified, communicated or responded during the same week, and during the same or following week. DSNO, Nord-Ouest department; DSN, Nord; DSNE, Nord-Est; DSA, Artibonite; DSC, Centre; DSO, Ouest; DSNi, Nippes; DSSE, Sud-Est; DSS, Sud; DSGA, Grand’Anse department.

### Response to cholera alerts by WaSH mobile teams

Between July 2013 and June 2017, WaSH mobile teams reported 31,306 field response interventions against cholera across the country, out of which 61% were conducted in communes in *red* alert and 14% in communes in *orange* alert (data no shown). The rest targeted *green* communes with sporadic cases (13%), with no case (7%), or commune with no data (6%).

Between July 2013 and June 2017, WaSH mobile teams responded to 49% of the 7,853 *red or orange* alerts during the same week, and to 59% of them during the same or following week (Table 3). Using generalized linear mixed models, response rates were significantly better for *red than* for *orange* alerts, whether during the week (OR, 2.52 [2.23–2.86]) or during the same or following week (Figure 3 Panel C, Table 3). The WaSH response rate to *red and orange* alerts significantly improved throughout the study period (Figure 3 Panel C, Table 3). For instance, response to *red* alerts during the same week climbed from 18% during 2013 2^n^ semester to 84% during 2017 1^st^ semester (Figure 3 Panel C). Rates for *orange* alerts were 10% and 67%, respectively, during the same periods (Figure 3 Panel C). *Red* alerts received a median of 1 response intervention during the same week [Interquartile range IQR, 0–5], which was significantly more than *orange* alerts (Relative risk (RR), 2.39 [2.20–2.59]) (Figure 4 Panel A, Table 4). The number of response interventions *per red or orange* alert significantly increased during the study, and it exhibited a significant heterogeneity between the 10 departments (*P*-value < 0.0001) (Figure 4 Panel A, Table 4).

**Figure 4.**
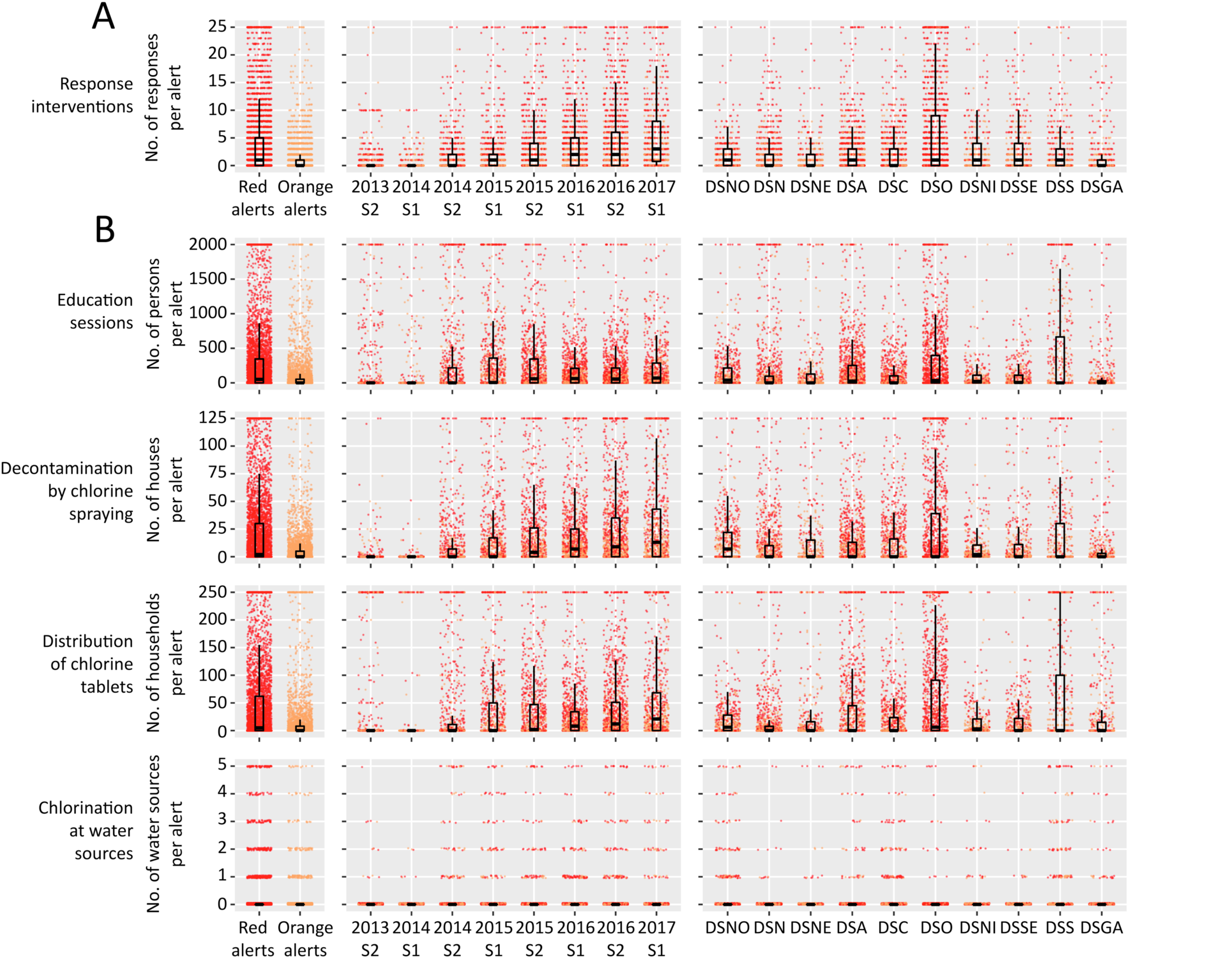
– Characteristics of WaSH response to cholera alerts from July 2013 to June 2017: number of response interventions (Panel A), number of people educated, number of decontaminated houses, number of households to which chlorine tablets were distributed and number of water sources that were chlorinated (Panel B) during the same week as alert; difference between alert levels (*red* vs *orange*), evolution along the eight semesters of the study period, and difference between departments. Dots show the characteristics of response to each *red* or *orange* alert. Boxplots show the median, interquartile range (IQR) and whiskers (1.5*IQR) of characteristics of response to alerts. DSNO, Nord-Ouest department; DSN, Nord; DSNE, Nord-Est; DSA, Artibonite; DSC, Centre; DSO, Ouest; DSNi, Nippes; DSSE, Sud-Est; DSS, Sud; DSGA, Grand’Anse department.

**Table 4.** – Characteristics of WaSH response to cholera alerts from July 2013 to June 2017: number of response interventions, number of people educated, number of decontaminated houses, number of households to which chlorine tablets were distributed and number of water sources that were chlorinated during the same week as alert; difference between alert levels (*red* vs *orange*), evolution along the eight semesters of the study period, and heterogeneity between departments.

| *Red or orange alerts*<br>(n = 7,853): | Median<br>[IQR*] | Difference<br>between alert levels<br>(*Red* vs *Orange* alerts) |  | Evolution<br>throughout the study<br>(per semester since July<br>2013) |  | Difference<br>between<br>departments<br>(vs DSO*) |
| --- | --- | --- | --- | --- | --- | --- |
|  |  | Relative risk<br>[95%-CI] | P-value | Relative risk<br>[95%-CI] | P-value | P-value |
| WaSH response during the same week, no.<br>of interventions per alert | 0<br>[0-3] | 2.39<br>[2.20-2.59] | < 0.0001 | 1.46<br>[1.43-1.48] | < 0.0001 | < 0.0001 |
| Education sessions during the same week,<br>no. of persons educated per alert | 0<br>[0-187] | 3.13<br>[2.66-3.69] | < 0.0001 | 1.08<br>[1.04-1.12] | < 0.001 | < 0.0001 |
| Decontamination by spraying during the<br>same week, no. of houses per alert | 0<br>[0-16] | 3.05<br>[2.70-3.44] | < 0.0001 | 1.77<br>[1.72-1.83] | < 0.0001 | < 0.0001 |
| Distribution of chlorine tablets during the<br>same week, no. of households per alert | 0<br>[0-32] | 2.80<br>[2.37-3.30] | < 0.0001 | 1.03<br>[1.02-1.04] | < 0.0001 | < 0.0001 |
| Chlorination at water sources during the<br>same week, no. of sources per alert | 0<br>[0-0] | 3.53<br>[2.68-4.65] | < 0.0001 | 1.28<br>[1.20-1.37] | < 0.0001 | < 0.0001 |

\* IQR, Interquartile range; DSO, West department

Overall, 47% of the 7,853 *red and orange* alerts received education sessions during the same week, which addressed a median of 209 persons per alert; 44% of alerts were responded by the decontamination of a median of 20 houses; chlorine tablets were distributed in 44% of alerts, to a median of 40 households; and chlorination at water sources was implemented in 7% of alerts (data not shown). The numbers of persons who were educated, of houses that were decontaminated, of households who received chlorine tablets and of water sources that were chlorinated during the same week were significantly higher for *red than* for *orange* alerts (Figure 4 Panel B, Table 4). These four indicators significantly increased during the study, and they were significantly heterogeneous between the 10 departments (Figure 4 Panel B, Table 4).

## Discussion

When the Haitian Government, UNICEF and their partners launched the cholera alert-response activities in July 2013, Haiti remained the most cholera-affected country worldwide [10] but few interventions aiming to prevent transmission were continued in the field [11]. For the first time since the beginning of the epidemic, a nationwide coordinated program aimed to mitigate cholera transmission by rapidly responding to every cholera suspected case through WaSH field interventions targeting their neighboring community. Data of 31,306 WaSH response interventions gathered and analyzed until June 2017 show that execution of such a strategy was feasible and its cost was less than USD 8 million per year.

Indeed, implementation of the cholera response strategy was initially laborious and remained heterogeneous across Haitian departments, because it was impeded by logistic obstacles when aiming to reach remote and mountainous localities, the heterogeneity of contracted organizations, and difficulties to coordinate their actions, notably with peripheral health authorities. Over several periods, it proved difficult to secure the funding of the strategy. This generated short term contracts, which likely impaired the response capacities and efficiency of mobile teams, like during the 1^st^ semester of 2014. During this period, response teams were also temporally weakened by administrative constraints and a severe but under-reported chikungunya epidemic that affected a large proportion of NGO staff and field workers [55]. Nevertheless, response interventions drastically improved in promptness, exhaustiveness and intensity over the study period, so that during 1^st^ semester 2017, 75% of *red* and *orange* alerts were responded within the same epidemiological week, a median of 70 persons were educated per *red* or *orange* alert, a median of 13 houses were decontaminated and a median of 21 households received water chlorination tablets.

This study also presented the original cholera alert system which was set by DELR to monitor the epidemic and the response strategy, and soon became popular at central and departmental levels. However, these *red* orange and *green* alerts presented important limits. The selected alert definitions could not be compared to any gold standard, and the 58% culture positivity ratio of *red* alerts suggests a background noise of non-relevant alerts which was probably due to non-choleric acute watery diarrhea cases seeking care in the largest or most advanced cholera treatment centers [5]. Conversely, the severity of outbreaks was not well captured by *red* alerts that represented a wide range of epidemic situations. Surveillance data used for alert identification did not include community cases, and the community deaths were inconstantly reported [5]. Therefore, some cholera outbreaks could have been overlooked, especially in remote areas. In addition, DELR prospectively communicated only 35% of cholera alerts during the same week, mainly because surveillance data were obtained with chronic delays. Communes in alert did not locate the actual patients’ place of residence but their treatment institution that notified cases. The weekly time scale of alerts was also too long compared with cholera median incubation of 1.4 days [56]. For these reasons, alerts were hardly used to rapidly guide field response interventions. Instead, mobile teams directly gathered their own epidemiological data in treatment institutions, and peripheral health authorities progressively established and shared detailed line listings of suspected cholera cases, which included patient address. Hence, response rates by WaSH mobile teams soon became higher than communication rates by DELR. Anyhow, identification and communication of alerts was progressively abandoned by DELR after departure of the epidemiologist in charge of this work.

To monitor the implementation of the alert-response strategy, we chose to use alerts that were retrospectively computed based on consolidated surveillance databases. These alerts proved a practical and original indicator, but they brought several limits to our analyses. Their weekly time scale largely exceeded the 48h response deadline that mobile teams were requested to respect, which may have overestimated response rates. Interventions were linked with alerts, based on the commune where patients were treated, whereas they were conducted locally, at patients’ homes and neighborhoods. Computed response rates were thus a surrogate of real response activities, which we believe relevant enough to assess the dynamic of the implementation of this strategy. A new reporting form of response interventions based on the line-listing has been used since the beginning of 2017, which should provide more accurate indicators for future analyses. The quality of reporting appeared heterogeneous between the organizations implementing response interventions, especially for exact quantities of distributed items we thus could not include in our analyses. Thanks to several evolutions of reporting forms, this improved along the study period. Unfortunately, we could not include response activities conducted by EMIRAs and medical organizations in our evaluation. Many of their interventions were common with reporting WaSH mobile teams, but available data about medical activities such as active case finding and chemoprophylaxis in the community was scarce. Moreover, several other organizations operated in community prevention of cholera during the study period, such as Brigada Médica Cubana, Médecins sans Frontières – Netherlands, Gheskio, Zanmi Lasante, Canadian Red Cross…but their field response activities to cholera cases appeared limited in comparison with the 31,306 WaSH response interventions included in the present study.

Additional evaluations are needed to better examine the quality of response interventions in terms of timing and geographic targeting, number of reached persons, methodology of education sessions, quantity of distributed water treatment products, as well as their impact on hand washing, defecation or water treatment practices [57,58]. Post-distribution monitoring has been encouraged from 2017 and their data should be analyzed. Considering the potential risk of resistance selection [49], use of chemoprophylaxis using doxycycline should be quantified and evaluated. After three years of reactive use, it seems that no resistant *V. cholerae* clinical strain has been isolated in Haiti so far (data not shown).

Evaluation of the efficiency and impact of this response strategy was out of this paper’s scope. This is a complex task and several methodological approaches are being considered. But together with other prevention activities conducted by MSPP, DINEPA and other organizations, the few oral vaccination campaigns implemented since 2012 [59], and the slow progress achieved in water infrastructure provision, this national alert-response strategy may have played a key role in the drops in cholera incidence observed in 2014 and in 2017. Unlike in 2014, late 2017 epidemiological reports even showed an unprecedented low cholera incidence in spite of the rainy season [3]. These achievements seemed unlikely according to the most recent and fitted cholera transmission model [60], PAHO predictions [61,62] and objectives of the 2013–2022 Cholera Elimination Plan [12].

The rapid response strategy still constitutes a core element of the 2016–2018 mid-term development of the national plan for cholera elimination of the Haitian Government [63], and of the new United Nations approach to cholera in Haiti that was adopted by the General Assembly in December 2016 [64]. However, it has become more and more difficult to get funding, and continuation of the strategy throughout 2018 is not guaranteed. It is mandatory to optimize future elimination efforts and thus to further evaluate the impact of each component of the cholera response. Results of ongoing dedicated studies will be very informative for actors involved in the implementation of cholera control strategies as well as international donors.

## Acknowledgements

We are grateful to the people who organized the strategy at the national and departmental levels. We also thank the staff of MSPP, UNICEF DINEPA and NGOs, who cared for patients, conducted alert investigations, implemented and coordinated field responses, gathered epidemiological and intervention data, analyzed stool cultures, or compiled and prospectively analyzed the cholera and interventions databases.

## Supplementary data

**Supplementary Figure S1.**
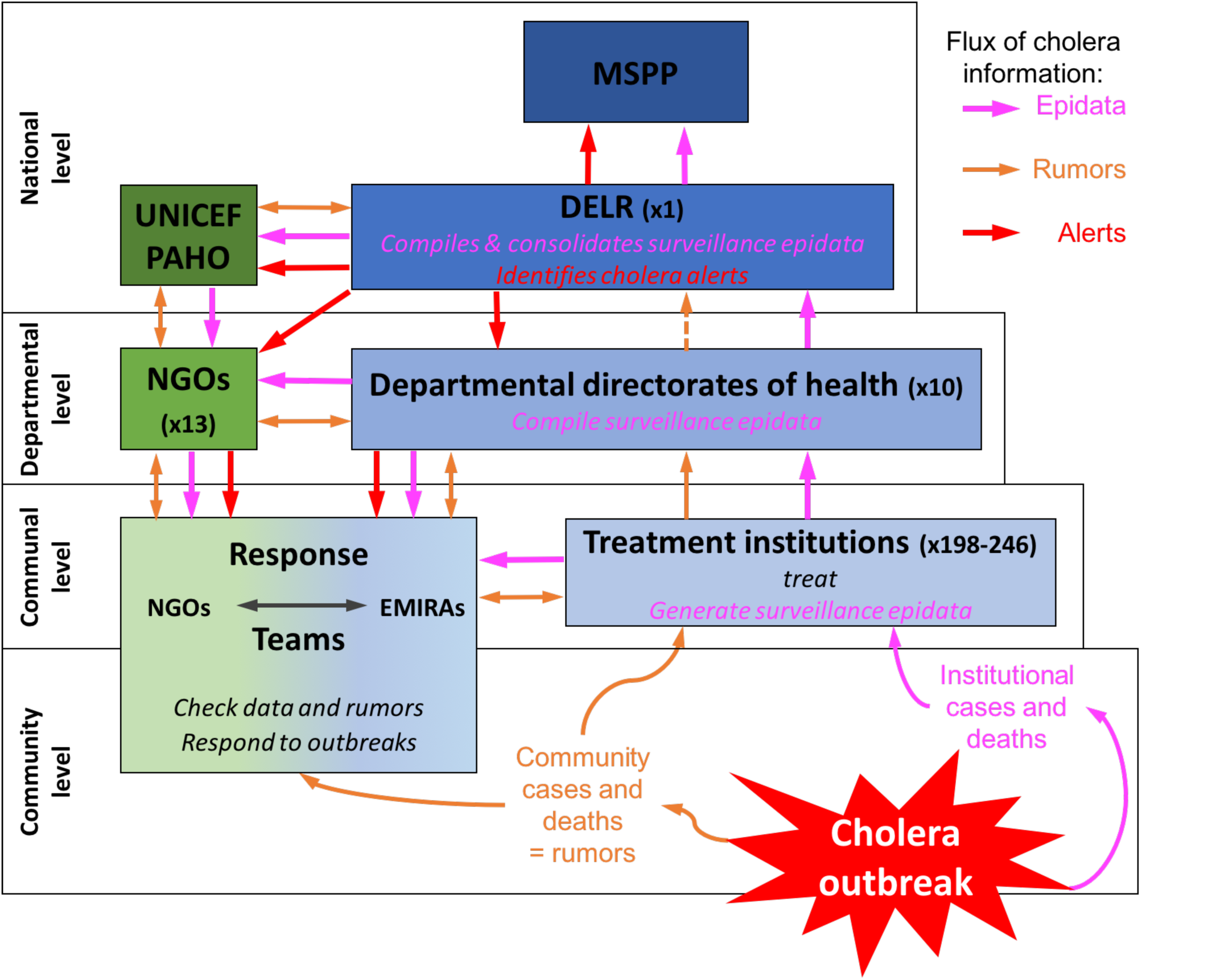
– Organization of cholera surveillance in Haiti.

**Supplementary Figure S2.**
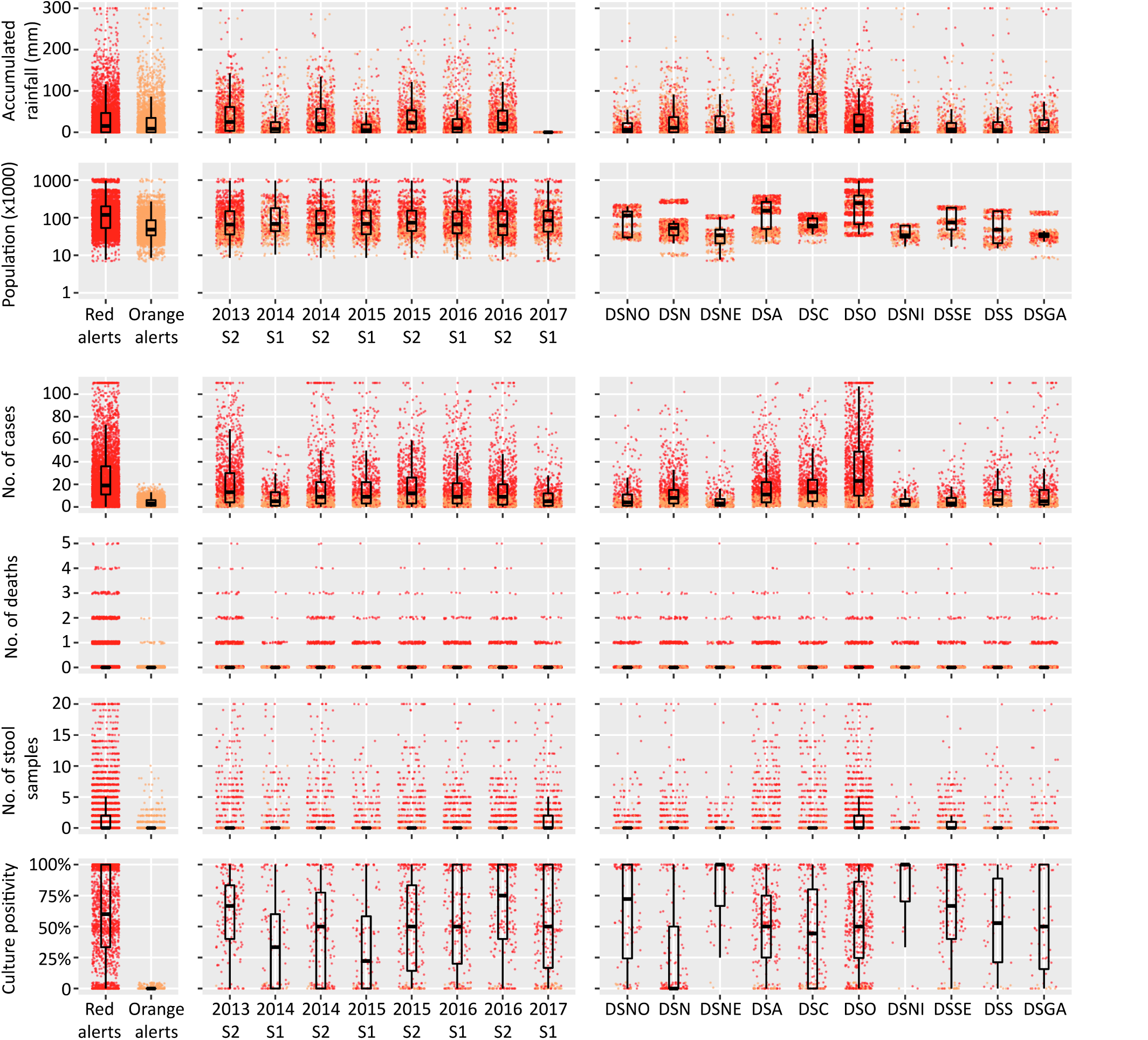
– Characteristics of retrospectively identified *red* and *orange* alerts from July 2013 to June 2017: difference between alert levels, evolution along the eight semesters of the study period, and difference between departments. Dots show the characteristics of each *red* or *orange* alert. Boxplots show the median, interquartile range (IQR) and whiskers (1.5*IQR) of alert characteristics. DSNO, Nord-Ouest department; DSN, Nord; DSNE, Nord-Est; DSA, Artibonite; DSC, Centre; DSO, Ouest; DSNi, Nippes; DSSE, Sud-Est; DSS, Sud; DSGA, Grand’Anse department.

**Supplementary Table S1.**
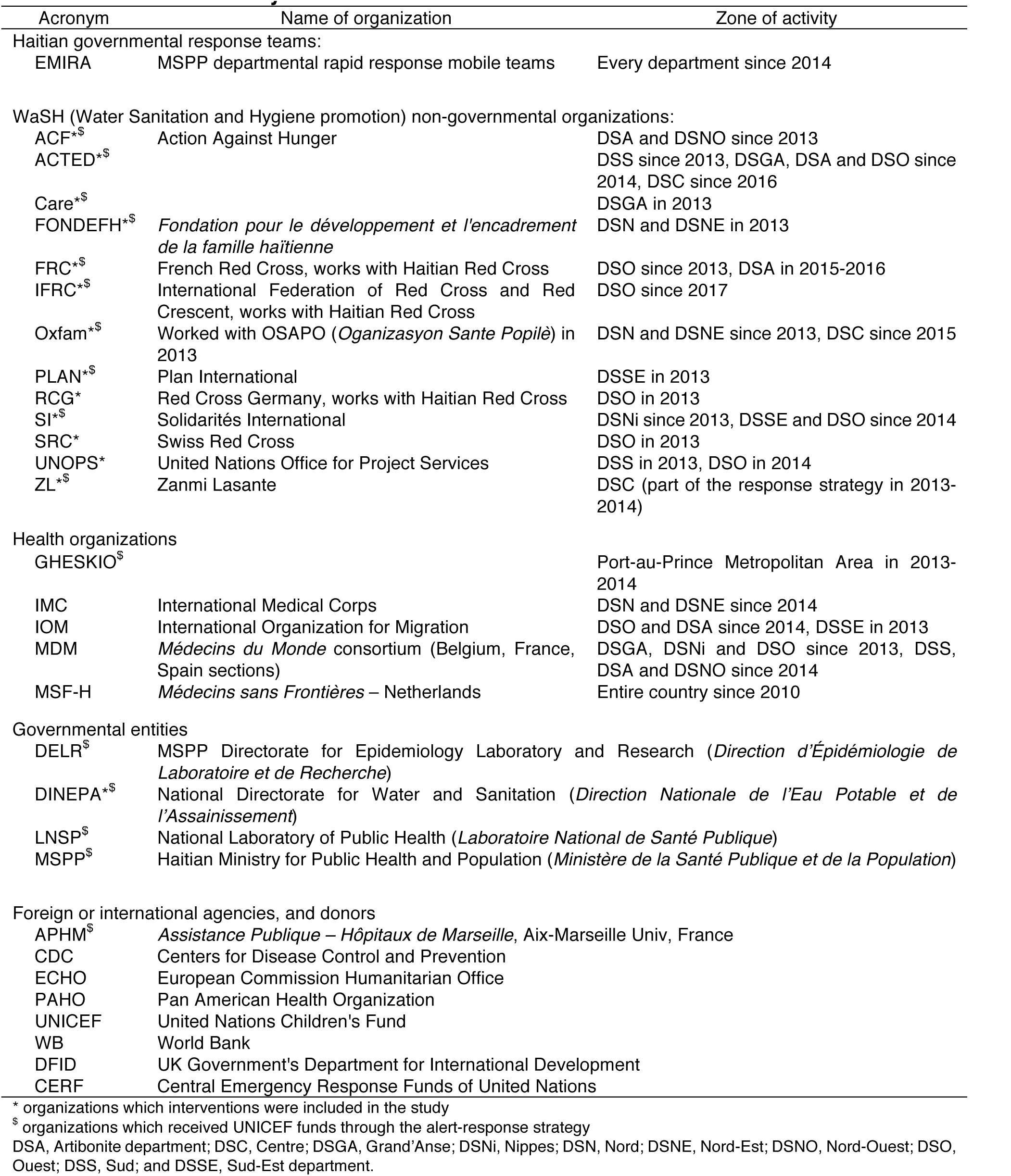
– Main actors involved in community response to cholera alerts and main areas of activity.

**Supplementary Table S2.** – Databases used in the study

| Database | Rows unit | Main variables of interest | Spatial scale | Time scale | Begin Date | End date | Database constitution | Data sources | Cleaning of data | Availability of database | Main potential sources of bias | Efforts to address potential sources of bias |
| --- | --- | --- | --- | --- | --- | --- | --- | --- | --- | --- | --- | --- |
| National cholera institutional database | Cholera treatment institutions | Eight variables: cumulated number of suspected cholera cases seen in treatment institutions or hospitalized, deaths in treatment institution and deaths in the community; each variable available for < 5 years and ≥ 5 years of age | Communes | Days | 10/18/2010 | ongoing | DELR | Cholera treatment institutions via departmental health directorates | Day to day prospective cleaning and quality-control using field information. Missing data for open institutions were set to zero | Upon approval by Haiti MSPP | Under-reporting, especially of community cholera cases and deaths. Low specificity of clinical definition | Prospective efforts to improve the sensitivity of the surveillance system. Promotion of stool sample *V. cholerae* cultures |
| Cholera treatment institution databases | Cholera treatment institutions | Institution opening and closing | Communes | Weeks | 07/30/2012 | Ongoing | Authors | Departmental health directorates, DELR, field investigations |  | Upon approval by Haiti MSPP | Misreporting of institutions opening and closing | Regular prospective field investigations |
| LNSP database | Stool samples | Date and place of sampling, date of sample reception by LNSP, result of *V. cholerae* O1 culture | Communes | Days | 10/19/2010 | Ongoing | LNSP | LNSP | No need | Upon approval by Haiti MSPP | No systematic stool sampling for *V. cholerae* culture. Decreased sensitivity if suboptimal transport conditions of samples | Prospective efforts to improve the exhaustiveness and quality of the cholera bacteriological surveillance system |
| Alert bulletin database | Cholera bulletins | Date of alert identification, date of bulletin diffusion, name of communes in alert | Communes | Days | 07/01/2013 | Ongoing | Authors | DELR | No need | Upon approval by Haiti MSPP | No |  |
| Intervention database | Field interventions | Names of mobile teams, place and date of intervention, conclusion of investigation, number of households decontaminated, number of people targeted by awareness campaigns, number of people receiving water treatment products, number of water chlorination points | Communes, communal sections, localities | Days | 07/01/2013 | Ongoing | UNICEF-Haiti | Response partners | Regular prospective cleaning and quality-control using field information. Missing data was set to zero | Upon approval by UNICEF-Haiti | Underreporting and misreporting of response interventions | Regular field investigations |
| Commune characteristic database | Communes | Coordinates, total population estimated in 2015, urban population estimated in 2015, coastal location, date of past OCV campaign | Communes | NA | NA | NA | Authors | IHSI*, MSPP | No need | Upon approval by corresponding author | Poor population estimates (no census since 2003) |  |
| Country and communal accumulated rainfall | Haiti and communes | Satellite-estimated daily accumulated rainfall | Communes | Days | 09/01/2010 | Ongoing | Authors | NASA websites<sup>\$</sup> | No need | Open source | Satellite remote sensing and modelling with a 0.25 degree resolution. No detection of potentially heavy local rainfall | |

DELR, Directorate of Epidemiology Laboratory and Research (MSPP); LNSP, National Laboratory of Public Health (MSPP); MSPP, Haiti Ministry of Public Health and Population; IHSI, Haitian Institute of Statistics and Informatics; NASA, US National Aeronautics and Space Administration; OCV, oral cholera vaccine
\* http://www.ihsi.ht/pdf/projection/Estimat_PopTotal_18ans_Menag2015.pdf (accessed 26 January 2018)
\$ http://giovanni.gsfc.nasa.gov/giovanni/ (accessed 26 January, 2018)

